# Differences in Relevant Physicochemical Properties Correlate with Synergistic Activity of Antimicrobial Peptides

**DOI:** 10.1101/2023.06.27.546817

**Authors:** Angela Medvedeva, Hamid Teimouri, Anatoly B. Kolomeisky

## Abstract

With the urgent need for new medical approaches due to increased bacterial resistance to antibiotics, antimicrobial peptides (AMPs) have been considered as potential treatments for infections. Experiments indicate that combinations of several types of AMPs might be more effective at inhibiting bacterial growth with reduced toxicity and a lower likelihood of inducing bacteria resistance. The molecular mechanisms of AMP-AMP synergistic antimicrobial activity, however, remain not well understood. Here, we present a theoretical approach that allows us to relate the physicochemical properties of AMPs and their antimicrobial cooperativity. A concept of physicochemical similarity is introduced, and it is found that less similar AMPs with respect to certain physicochemical properties lead to greater synergy because of their complementary antibacterial actions. The analysis of correlations between the similarity and the antimicrobial properties allows us to effectively separate synergistic from non-synergistic AMPs pairs. Our theoretical approach can be used for the rational design of more effective AMPs combinations for specific bacterial targets, for clarifying the mechanisms of bacterial elimination, and for a better understanding of cooperativity phenomena in biological systems.

**Author summary:** It is impossible to imagine modern medicine without antibiotics. But there is a growing problem of increased bacterial resistance to them. These considerations stimulated a search for novel methods to defend against infections. Antimicrobial peptides (AMPs) came out as powerful antibacterial agents. It was also found that combinations of AMPs are even more efficient than individual peptides. The mechanisms of such synergistic activities, however, are not understood. We developed a computational framework that allows us to connect the physicochemical properties of AMPs and their abilities to cooperatively eliminate infections. It is found that less similar peptides might exhibit synergy because of their complementary antibacterial properties. Our theoretical approach might lead to a better rational design of new antimicrobial drugs.

## Introduction

One of the main achievements of modern medicine is the ability to efficiently eliminate various infections. This is currently done by using several classes of specific small organic molecules that are generally called antibiotics. But in the last 30 years, we are witnessing an increasing resistance to antibiotics in bacteria which threatens to severely decrease our ability to protect human health [1–3]. These alarming facts stimulated a broad search for novel anti-bacterial agents and techniques. The anti-microbial peptides (AMPs), which are produced by multi-cellular organisms as part of their immune responses to external infections, came out as promising alternatives to antibiotics [4–9]. AMPs are relatively short peptide-chain molecules with large fractions of separated positively charged and hydrophobic residues that exhibit activities against multiple classes of bacteria, fungi, viruses, and even cancer [10–14]. It was also observed that combinations of some specific types of AMPs frequently work much more efficiently than single-type peptides [15–19]. Although it is known that AMPs combinations are less toxic, can hinder the ability of bacteria to develop resistance, and can associate to bacterial cells faster, the mechanisms of AMPs cooperativity remain not well understood [18–21].

AMPs exhibit a wide spectrum of structures and mechanisms of bacterial removal [5, 6]. But the dominating antimicrobial pathway is the association of AMPs to bacterial membranes with the following pore formation that leads to the death of the bacterial cell [12, 22, 23]. The efficiency of antimicrobial peptides in eliminating infections is measured by the minimal inhibitory concentration (MIC) required to inhibit the growth of the bacterial population. It is interesting that some AMPs might also sensitize antibiotics in their action against the previously-resistant bacterium [6]. In addition, like antibiotics, some AMPs have a broad spectrum of antimicrobial activity, targeting multiple different species of bacteria. But unlike antibiotics, AMP-based drugs are powerful against antibiotic-resistant strains. The combinations of different types of AMPs might be even more efficient in their antibacterial activities [16, 17, 19]. They show higher efficacy, reduced toxicity, and a lower likelihood of inducing bacteria resistance compared to available antibiotics and even individual AMPs. The efficiency of AMP-AMP combinations is measured by fractional inhibitory concentrations (FIC), which reflect the extent to which MIC for a given AMP in combination with another AMP is reduced compared to the MIC of the same AMP when applied individually.

The microscopic origin of antibacterial cooperativity between different types of AMPs is unclear. Bacterial resistance to individual AMPs is rare yet possible [24–27], while bacterial resistance is less probable to AMP-AMP combinations [18, 28]. This is one of the reasons why AMP-AMP combinations are more likely to not only be more efficient than individual AMPs [29–31] but also serve as long-term antimicrobials in contrast to traditional antibiotics [32–34]. Several other possible sources of cooperativity have been proposed. It was suggested, based on experimental findings, that two AMPs might be synergistic because they individually target different types of bacteria [e.g., gram-positive and gram-negative [35]]. Thus, their combinations are more effective against both types of bacteria [36]. Furthermore, two AMPs may have different mechanisms of action to disrupt the bacterial function [37–41], or they might perform different antimicrobial functions (e.g., one AMP sensitizes the bacterium cell to the other AMP). Then, the existence of multiple antimicrobial mechanisms can prevent the development of resistance to the combination since bacteria are not able to simultaneously respond to both of them [42]. Also, two AMPs with different secondary structures (for example, *α* − helix and *β*-sheet) might also better inhibit bacterial growth [43–46], and the difference in their structures can prevent the peptide aggregation that should decrease the antibacterial properties.

It is clear that possible synergy between two different types of AMPs is a result of direct or indirect molecular interactions. It was suggested that this interaction might appear due to similarity in the structures of the peptides [47]. However, recent experimental studies of individual AMPs determined that other physicochemical properties, and not structures, correlate better with higher antimicrobial activity and a lower likelihood of bacteria resistance [48, 49]. In addition, for two AMPs to be a synergistic pair it was shown that they must have very different hydrophobicities [38, 44, 50]. Furthermore, it was proposed that the larger differences in physicochemical properties for AMPs combinations might be needed to prevent bacterial resistance. This is because bacteria are less likely to transfer resistance from an AMP in one class characterized by certain physicochemical features to an AMP in a different class (characterized by different physicochemical features than the first class), a phenomenon known as cross-resistance [51–53]. But experimental tests of this idea exhibit mixed results. Synergy and lack of cross-resistance are evident in the example of food-preservative AMPs curvaticin-13 and nisin [54], while AMPs microcins showed cross-resistance and lack of synergistic antimicrobial activity [55].

In this paper, we postulate that two classes of AMPs will cooperate if they have very different physicochemical properties relevant to the antimicrobial activity because they will complement each other. To test this hypothesis, we introduce a concept of physicochemical similarity defined as Euclidean distance in the space of more than 1500 physicochemical descriptors. This quantitative approach explicitly evaluates the correlations between physicochemical similarities and FICs of different AMP-AMP combinations. This allows us to select the most important features that contribute to synergistic antimicrobial activity. Applying Principal Component Analysis (PCA), it is found that it is possible to distinguish between effective and ineffective AMP-AMP combinations only when similarity is calculated in terms of the selected features. We argue that greater physicochemical dissimilarity between AMPs in certain features is associated with stronger cooperative antimicrobial activity. Possible mesoscopic arguments to support these observations are presented. Our computational approach can be used to rationalize the design of effective and bacteria-specific AMP-AMP combinations as potential drugs, and it can also assist in clarifying more microscopic picture of the bacterial removal by AMPs.

## Results

### Feature selection based on correlations between FIC values and physicochemical similarity of AMP pairs

The quantitative measure of the efficiency of single-type AMPs against a bacterial species is MIC, which is the minimum concentration of an antimicrobial agent required to completely inhibit the growth of a bacterium population. The corresponding quantitative measure for AMP-AMP combinations is FIC, which is the sum of the ratios of each peptide’s MIC in the combination to individual MIC,

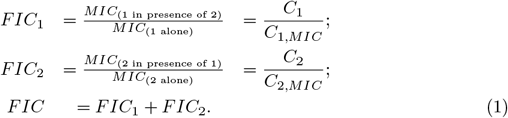

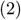

To better understand the cooperativity of two types of AMPs, it is convenient to look into a schematic view of a checkboard assay, shown in Fig. 1, which is typically utilized in experiments on measuring synergy of AMPs [56]. In this graph, the concentrations of peptides are presented as fractions of their corresponding MICs for single-type AMP measurements, and each circle corresponds to a specific combination of AMPs. Different curves describe the conditions at which bacterial growth is stopped.

**Fig 1.**
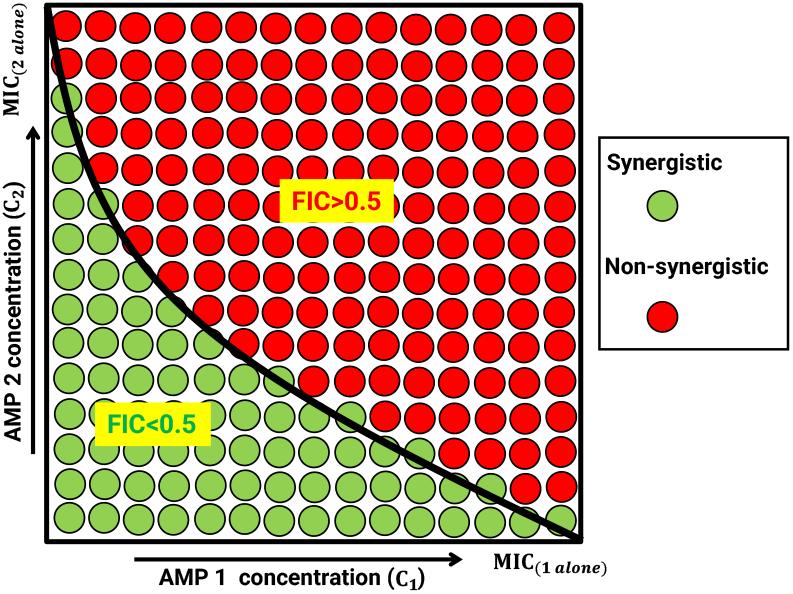
A schematic view of the checkerboard assay in which AMP-AMP combinations are tested. Each circle represents different sets of concentrations of AMPs. The solid black curve describes the conditions at which bacterial growth stops.

For all curves below the black curve (e.g., green circles), we have *FIC* < 0.5, and this describes the synergistic action of AMP pairs in the elimination of bacterial infection. The presence of the second type of AMP enhances the antibacterial efficiency of the first type of AMP. For all curves above the black curve (red circles), we have *FIC* > 0.5, and these conditions are viewed in our method as non-synergistic for AMPs combinations. In other words, the presence of the second type of AMP lowers or does not affect the antimicrobial efficiency of the first type of AMP.

For practical purposes, one is always searching for strong synergistic pairs of AMPs, and then some arbitrary thresholds are utilized to define this region of antimicrobial activities. Moreover, to simplify the statistical analysis, it is convenient to have similar numbers of synergistic and non-synergistic pairs. In our work, *FIC* = 0.47 is chosen as the threshold for synergistic pairs, and AMP-AMP pairs with *FIC* > 0.47 are considered as effectively non-synergistic, and pairs with *FIC* < 0.47 are viewed as synergistic. We collapsed across weakly-synergistic, additive, and antagonistic categories because there was not enough data to examine each case separately, and our focus is on strongly synergistic pairs [57, 58]. It is important to note that the exact choice of the threshold value does not affect our main conclusions.

To obtain a quantitative description of the antibacterial efficiency of AMPs pairs, the data were extracted from the DBAASP database [59], and we ensured that the data we included in the analysis were collected under the same experimental conditions. These data contain FIC values for AMP-AMP combinations for three different bacterial species (*E. coli, M. luteus*, and *P. aeruginosa*), and in which both peptides in the combination consisted of only natural amino acids with an overall sequence length of at least 11 (see Table. 1). The latter requirement is needed for the subsequent proper extraction of physicochemical descriptors of peptides.

**Table 1.** Overview of the datasets for AMP-AMP combinations extracted from DBAASP database. Specifically, the number of synergistic and non-synergistic AMP pairs for each bacterium is presented.

| Bacteria | Synergistic AMP-AMP combinations | Non-synergistic AMP-AMP combinations |
| --- | --- | --- |
| *E. coli* | 12 | 50 |
| *M. Luteus* | 6 | 12 |
| *P. Aeruginosa* | 4 | 11 |

**Table 2.**
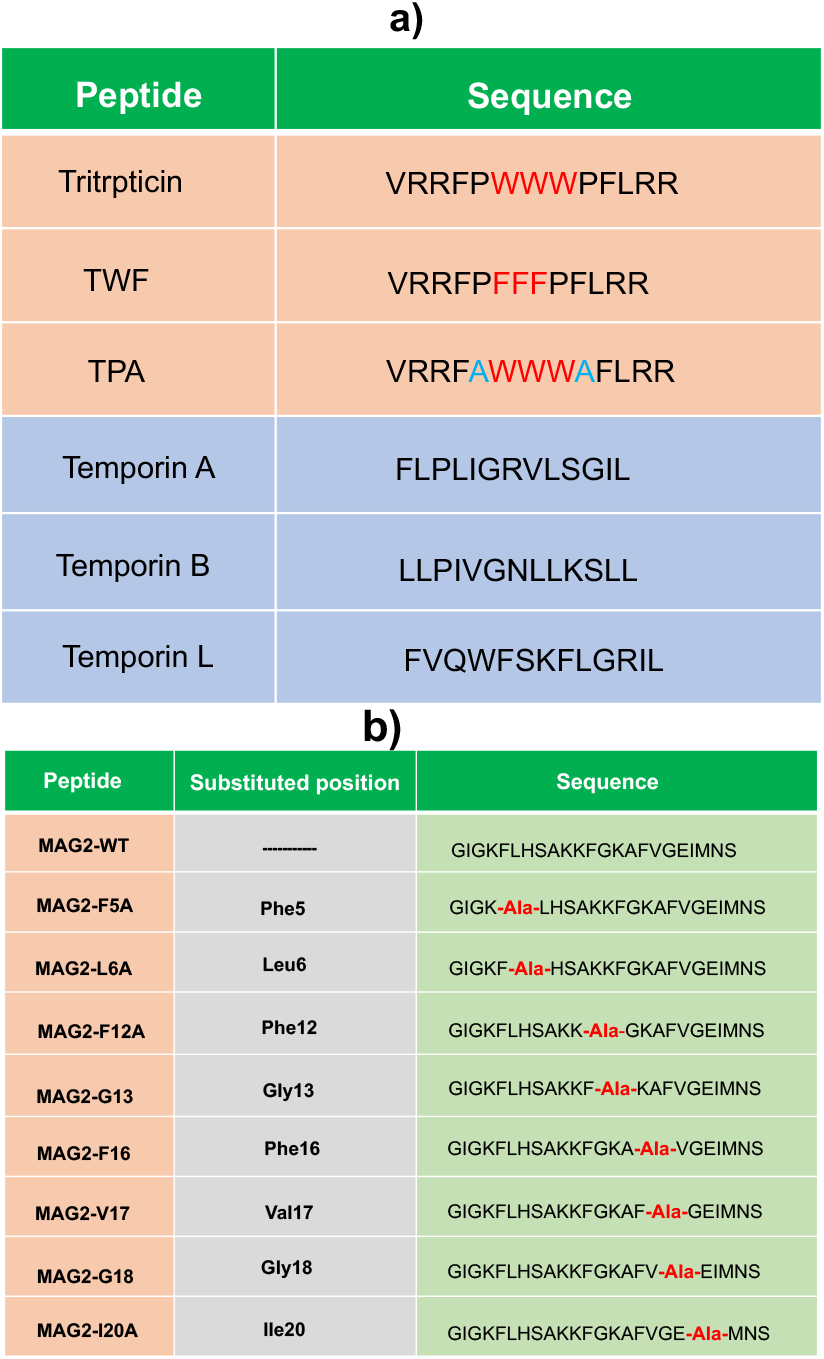
a) Various Tritrpticin and Temporin peptides and their corresponding sequences. b) Various Magainin-2 peptides and their corresponding sequences.

We extracted a comprehensive set of 1547 physicochemical properties for each peptide sequence using a bioinformatic package *propy* [60]. These physicochemical features include amino acid compositions (percentage of each amino acid in the peptide), net charge, hydrophobicity, polarizability, polarity, van der Waals forces, and solvent accessibility.

We first calculated the Euclidean distance, representing the inverse of the similarity of two peptides, between the AMPs in each pair in terms of all features to arrive at a single value [61, 62]. Then we calculated Spearman’s rank correlation coefficient [63, 64] between the Euclidean distances and FIC values for each AMP pair (see Materials and methods for details of the calculations).

Since there was no significant correlation between FIC and Euclidean distance for any pairs of peptides in terms of *all* descriptors, *p* > 0.05, we analyzed this relationship separately in terms of each of the 1547 physicochemical features. The corresponding histograms of correlation coefficients for three species of bacteria are shown in Fig. 2. We decided to choose only those features for which the Spearman correlation coefficient was statistically significant, i.e., |*r*_*s*_| is large. Furthermore, a statistical analysis with *p*-value of less than 0.005 was applied to filter irrelevant features. This corresponds to features with *r*_*S*_ smaller than red dashed lines for *r*_*S*_ < 0 and features with *r*_*S*_ larger than the red dashed lines for *r*_*s*_ > 0: see Fig. 2. Based on these criteria, we obtained the features shown in Fig. 3a for *E. coli*, in Fig. 3b for *M. luteus*, and in Fig. 3c for *P. aeruginosa*. Then, the physicochemical similarity for AMP pairs has been computed utilizing only the *selected* features to result in a single Euclidean distance value for each AMP pair, and the relationship between the FIC values and the distances has been analyzed (see Fig. 4). One can see that the distance in the space of selected physicochemical features between two peptides can be utilized to separate synergistic from non-synergistic AMP pairs.

**Fig 2.**
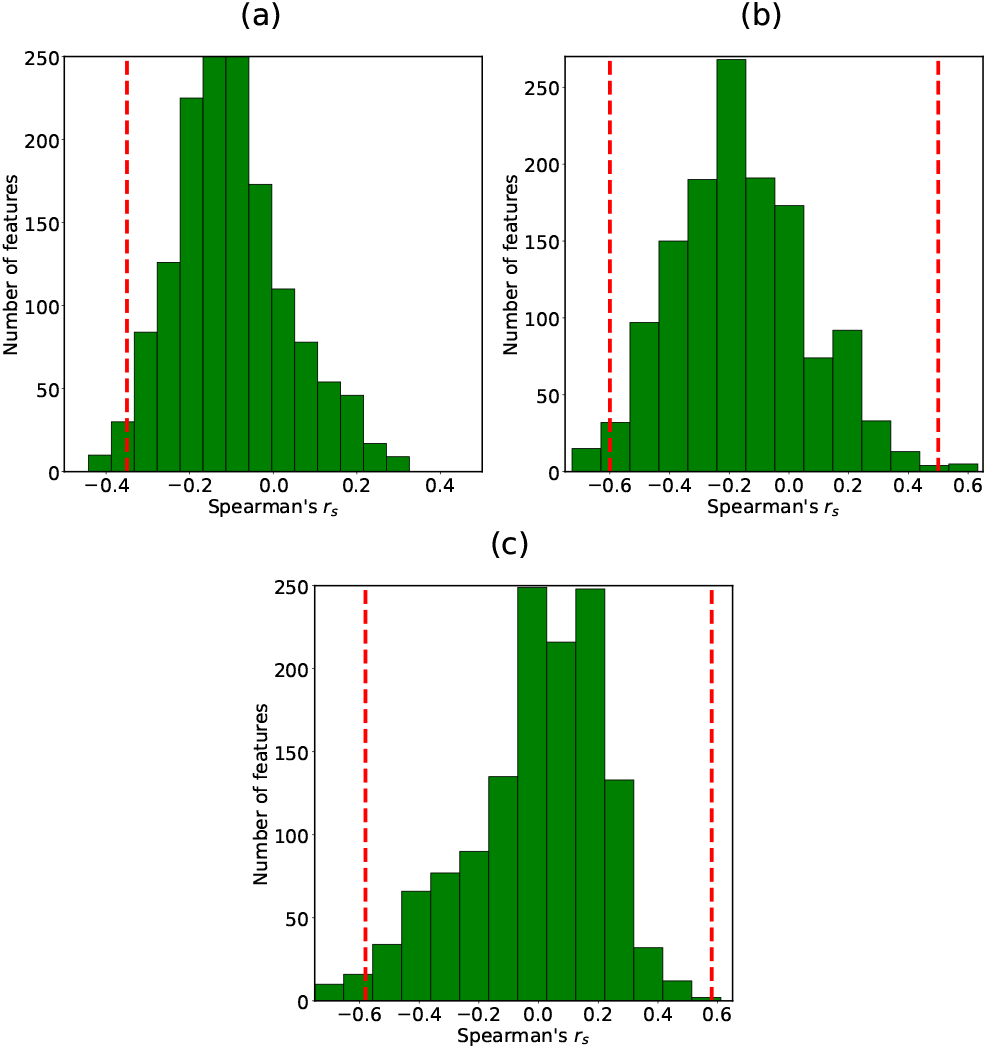
Distribution of Spearman correlation coefficients between fractional inhibitory concentration (FIC) values and the corresponding distances between paired AMPs in terms of individual physicochemical features (*d*^(Individual Feature)^(*A, B*)) for: a) *E. coli*, b) *M. luteus*, c) *P. aeruginosa*. Vertical red dashed lines show threshold values of Spearman’s correlations above which the corresponding features are selected.

**Fig 3.**
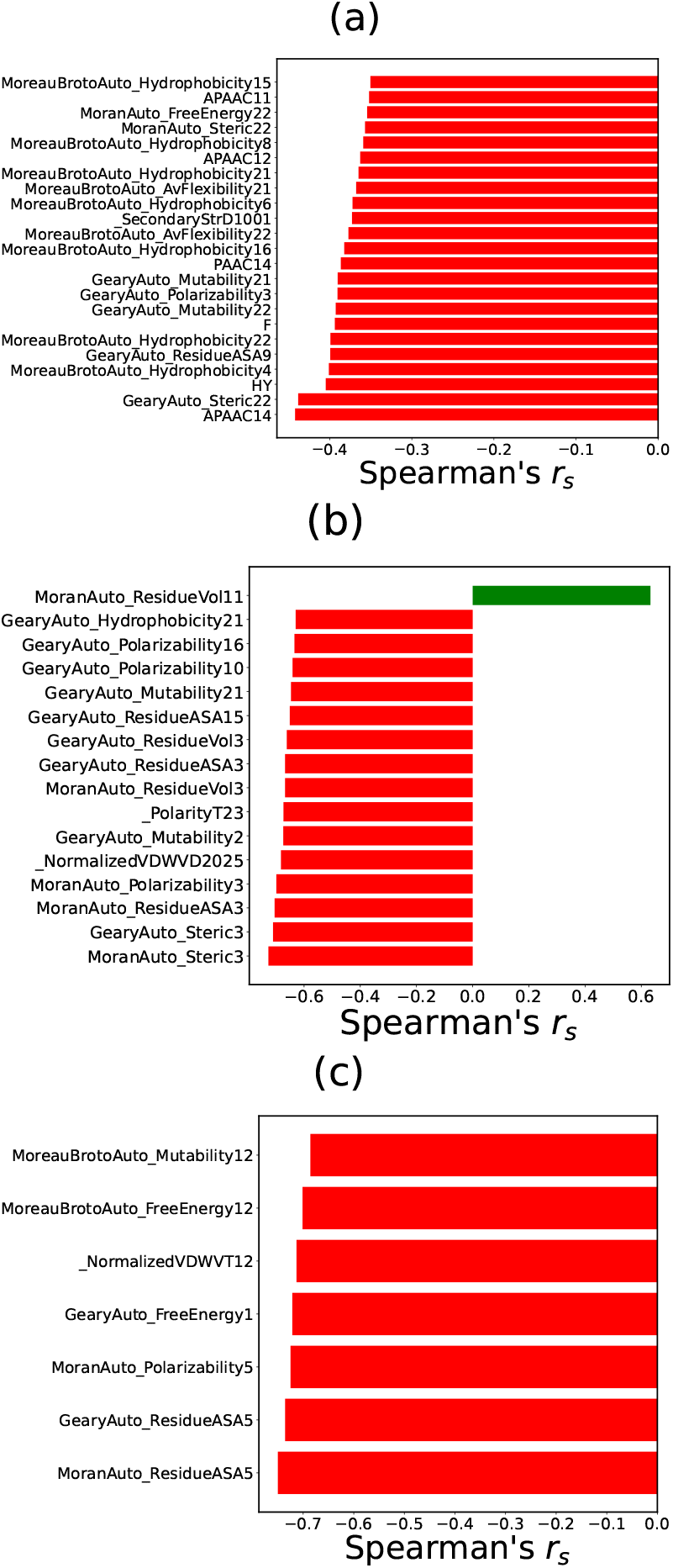
Relative importance of physicochemical features in terms of which AMP-AMP similarity is highly correlated with the corresponding FIC values: a) for *E. coli*, b) for *M. luteus*, c) for *P. aeruginosa*

**Fig 4.**
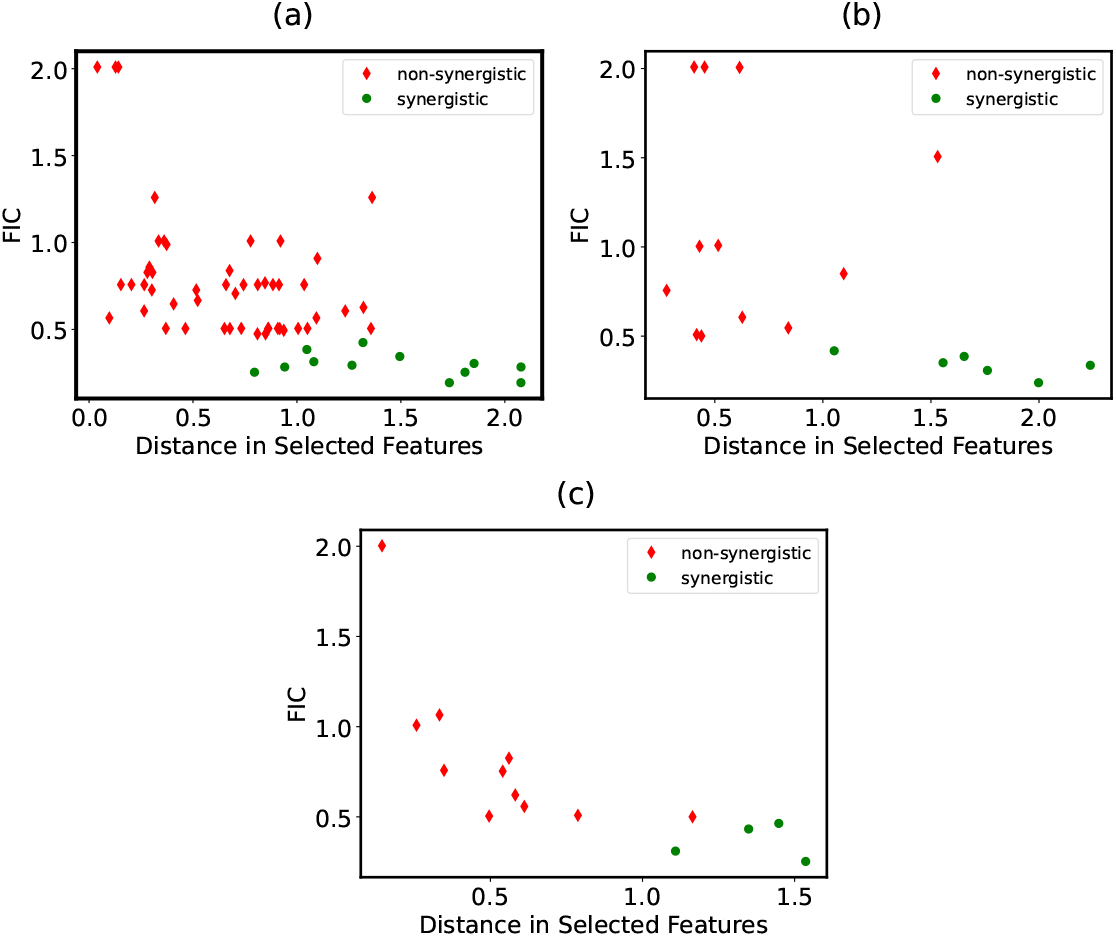
FIC versus distance in terms of selected features, as shown in Fig. 3 for AMP-AMP combinations targeting: a) *E. coli*, b)*M. luteus* c) *P. aeruginosa*

Our analysis suggests that most physicochemical features do not affect the correlations between the FIC and the similarity. The features that we selected are shown in Fig. 3a for *E. coli* bacterium. In Fig. 4a, for *E. coli* we plotted the distance between AMPs in the space of only selected features against the FIC values, and, in contrast to the case when all features are included (not shown), there is a visible separation between synergistic and non-synergistic AMP pairs. Similar results are observed for other analyzed bacterial species (see Figs. 4b and 4c). It is interesting to note that most selected physicochemical features are distinct for each bacterium (Fig. 3), but there was also overlap in certain properties, including hydrophobicity and polarizability, suggesting that there might be universal physicochemical features of AMPs that are important for eliminating any bacteria.

### Dimensionality reduction using selected features

Alternatively, various dimensionality reduction methods can be also explored to distinguish cooperating AMPs combinations from non-cooperative pairs. To further test the selected features in distinguishing synergistic from non-synergistic combinations, we opted to use principal component analysis (PCA) to map the data into a two-dimensional space in which synergistic pairs are distinguished from antagonistic pairs. To do so, we calculated AMP-AMP Euclidean distance in terms of each individual feature. Then each AMP-AMP pair could be represented as a point in *M* -dimensional space of the selected feature (see Materials and methods for more detail). The results of PCA confirmed that there is a separation between synergistic and non-synergistic combinations based on the selected features for each bacterium (see Fig. 5).

**Fig 5.**
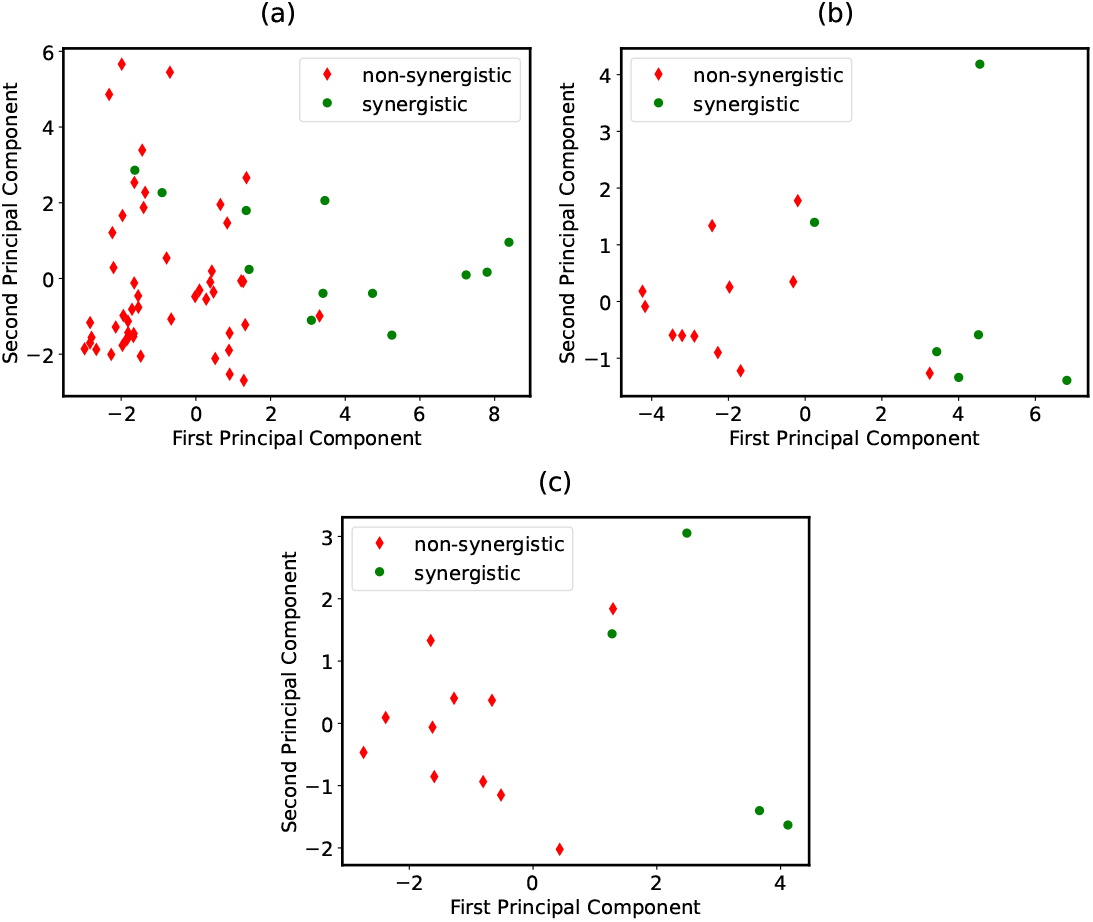
Principal component analysis analysis for separation of synergistic and non-synergistic AMP-AMP pairs targeting: a) *E. coli* b) *M. luteus* c) *P. aeruginosa*

### Feature selection based on correlations between FIC values and physicochemical similarity of dimers

Given the possibility of separating synergistic vs non-synergistic AMP-AMP pairs based on physicochemical similarity, we also explored the possible ways to distinguish antimicrobial vs non-antimicrobial AMP dimers. AMP dimers are molecules in which two AMPs are linked together by a chemical bond. As for AMP-AMP pairs, the information for AMP dimers was extracted from the DBAASP database, and since dimers act as one AMP, MIC values were collected instead of FIC values [59]. Moreover, the same criteria were applied to dimers as for AMP-AMP pairs: each sequence in the dimer must contain at least 11 amino acids, and the data must be collected under the same experimental conditions with the same bacterium as the target. The largest dataset of dimers that met the criteria was tested against *E. faecalis*. In this system, dimers with MIC values greater than or equal to 50 *μ*M are considered to be non-antimicrobial [65, 66]. Where there was more than one MIC value for the same dimer, we averaged and calculated the standard deviation. We removed dimers with averages corresponding to standard deviations greater than 0.2. Thus, we ensured based on the information in the DBAASP database that the 19 dimers in our dataset had reliable MIC values, i.e. that were consistent across experiments. The MIC values ranged from 0.0002 to 200.

As in the analysis of synergistic activity for combinations of AMPs, we calculated the Euclidean distance for each feature between the AMPs in each dimer. Then we correlated the Euclidean distances for each feature for each dimer with corresponding MICs for the dimers. The distribution of Spearman correlation coefficients is shown in Fig. 6a. We plotted the Euclidean distance for the feature set against MIC, shown in Fig 6b) for the statistically-significant features with the highest correlations, shown in Fig 6c). The most pronounced set of features is related to dipeptide composition, the percentage of a specific amino acid pair in the sequence, and this set of features is distinct from the features that were related to the synergistic antimicrobial activity for *E. coli, P. aeruginosa*, and *P. aeruginosa*, for which dipeptide composition was not a consistent feature. Notably, however, there was an overlap between the two datasets (AMP-AMP combinations vs dimers) in autocorrelation-related features.

**Fig 6.**
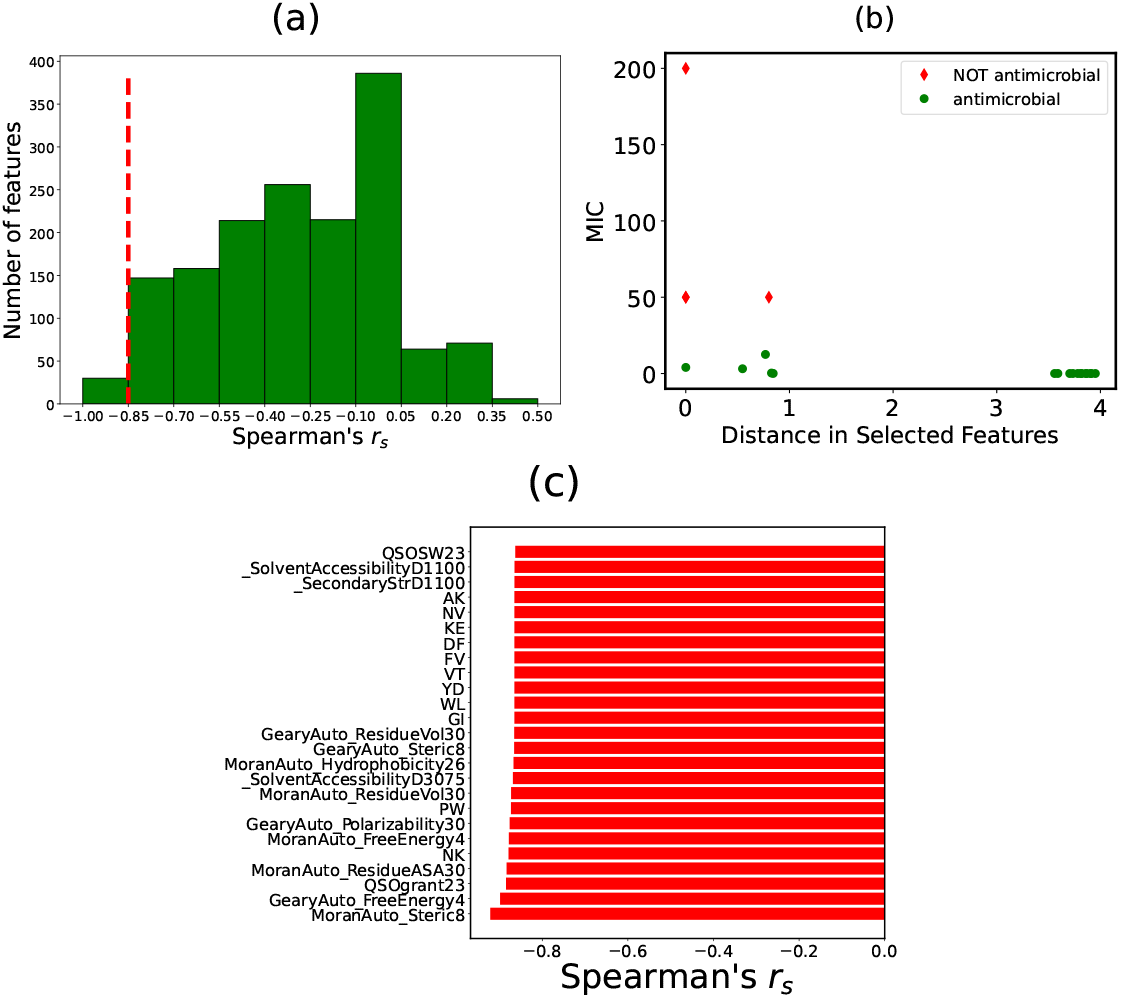
a) Distribution of Spearman correlation coefficients for all features for *E. faecalis* hetero-dimers b) Scatter plot of AMP-AMP distance versus MIC for hetero-dimers in terms of 25 statistically-significant selected features. c) Feature importance plot based Spearman correlation between MIC and AMP-AMP similarity of hetero-dimers in terms of individual features.

As shown in Fig 6b, the non-antimicrobial dimers in red are more similar in selected physicochemical characteristics alongside dimers with a certain spectrum of antimicrobial activity (*MIC* = 0.05*μM* - 12.5*μM*), whereas for the antimicrobial dimers with the lowest antimicrobial activity (*MIC <* 0.05*μM*), there were greater differences in selected features. This suggests that the strongest antimicrobial activity corresponds to larger differences in specific physicochemical features, including dipeptide composition and autocorrelation, between the two components of the dimer. Greater differences between the constituent monomers were associated with greater antimicrobial activity as reflected by lower MIC.

In summary, an analysis of the similarity of physicochemical features between components of each dimer revealed that highly antimicrobial vs non-antimicrobial dimers can be distinguished by their dissimilarity in specific physicochemical features that correlate with lower MIC, corresponding to higher antimicrobial activity. The findings are consistent with the analysis of the role of similarity in physicochemical features in the synergistic activity of AMP-AMP combinations; synergistic AMP-AMP combinations showed greater dissimilarity in the features that showed the strongest correlations with lower FIC and thus stronger antimicrobial effects.

### Analysis of specific selected features: autocorrelation functions

Our theoretical method indicates that among the important selected physicochemical features for all types of investigated bacteria are various autocorrelation functions, which measure the variation of different physicochemical properties for any pair of amino acid residues along the peptide sequence. There are three versions of auto-correlation functions, known as Geary, Moran, and Moreau-Broto [67, 68]. Although there is no fundamental difference between these functions, unlike Moreau-Broto, Geary and Moran auto-correlation parameters utilize averages and variances for each property. For example, the Moreau-Broto autocorrelation coefficient, which measures the correlation between physicochemical properties of residue *i* and residue *i* + *d* (along the peptide contour), is given by [68]

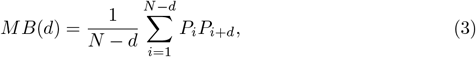

where *P*_*i*_ and *P*_*i*+*d*_ are physicochemical properties of residue *i* and residue *i* + *d*, respectively. Alternatively, we can define Geary autocorrelation function [68],

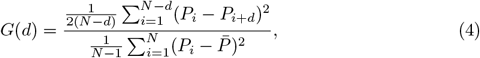

where 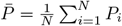 is the average physicochemical property of the sequence. Finally, one can define the Moran autocorrelation function, which is similar to Pearson’s correlation between the physicochemical property of residue *i* and residue *i* + *d* [68]:

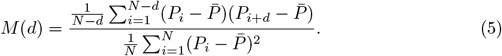

Considering physicochemical features important for *E. coli*, one can notice that dissimilarity of two AMPs in terms of autocorrelation in hydrophobicity contributes to synergistic activity of their combination. In this case, a positive autocorrelation corresponds to a peptide which is either totally hydrophobic or totally hydrophilic, i.e., it has the same sign of hydrophobicity for different amino acids along the peptide chain. A negative autocorrelation describes amphiphaticity when one side of the peptide is hydrophobic and the other side is hydrophilic, i.e., a change of sign in hydrophobicity along the peptide chain. Accordingly, a large difference in hydrophobicity autocorrelation coefficients between two peptides in a combination suggests that in the synergistic combination one peptide is most probably amphipathic, while the other is predominantly hydrophobic or hydrophilic.

To illustrate the idea that for two AMPs to cooperate in removing the bacteria they must be very different in terms of autocorrelations in hydrophobicity, let us analyze a specific example of AMP-AMP combinations that target *E*.*coli* bacteria. Three different types of peptides, Tritrpticin and its two derivatives labeled as TPA and TWF, are considered first [69]. Bacterial killing assays showed that the combination of Tritrpticin and TPA was not synergistic, while Tritrpticin-TWF and TPA-TWF pairs successfully cooperated against *E. coli*. To elucidate the relationship between the physicochemical similarity of these AMP pairs and their antibacterial activity, we computed the Moran autocorrelation parameter for hydrophobicity (which characterizes the degree of amphiphaticity of the peptide) at different distances along the peptide chain using the information from the *propy* package. The parameter *d* specifies the distance between amino acids along the peptide chain. As one can see in Fig. 7a, Tritrpticin is similar to TPA in terms of the lack of amphiphaticity, but it is different from TWF. Thus, the synergistic activity of TPA-TWF and Tritrpticin-TWF combinations might be attributed to the fact that TWF is highly amphiphatic in contrast to both Tritrpticin and TPA. At the same time, the amphiphaticity is absent in both Tritrpticin and TPA so that they are physicochemically more similar, making their combination ineffective against *E. coli*.

**Fig 7.**
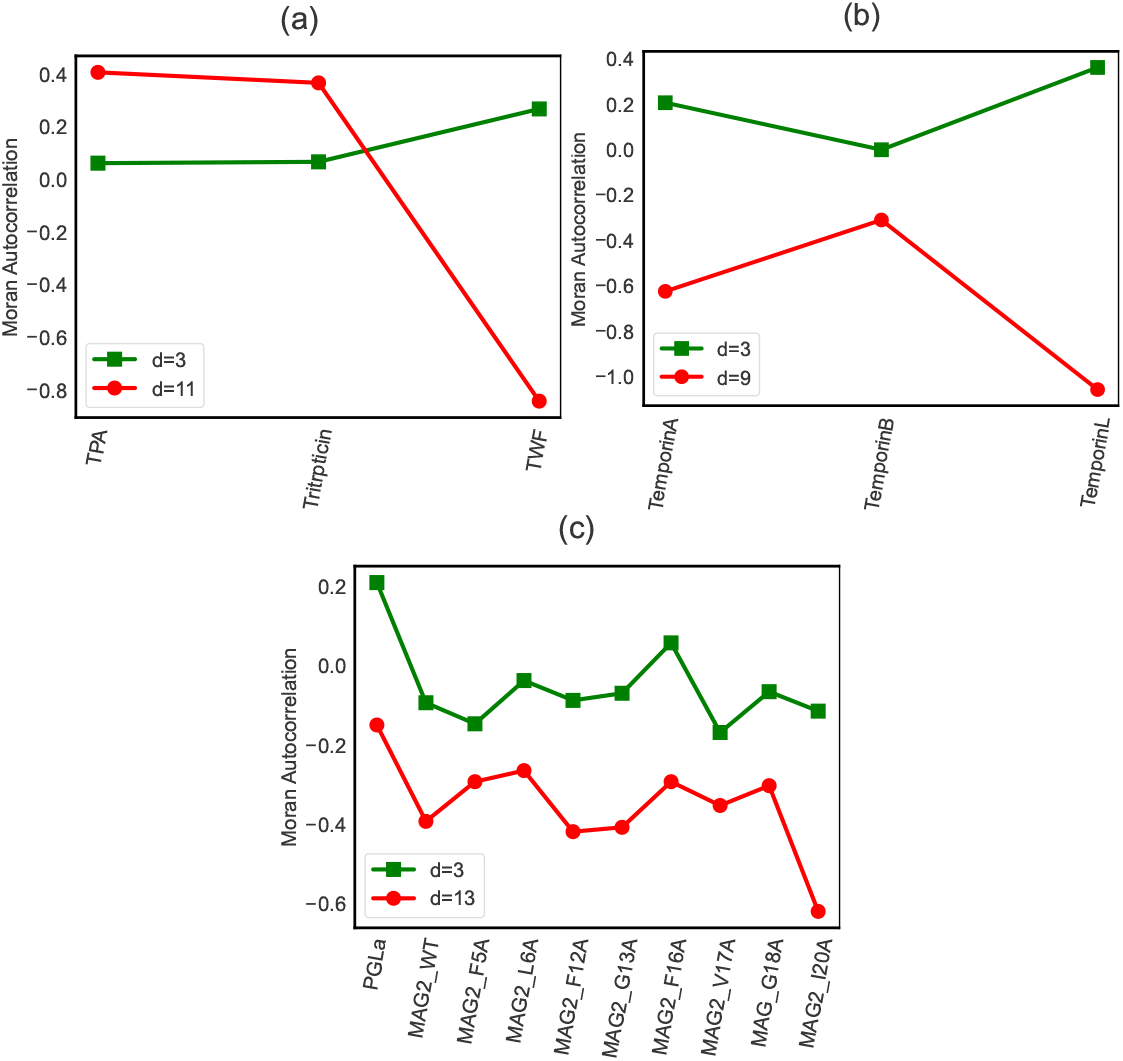
Calculation of Moran auto-correlation coefficient in hydrophobicity for a) Tritrpticin and its derivatives, TWF and TPA [69]. b) Temporins A, B, and L. Temporins A and B are non-synergistic, while Temporin L is synergistic with Temporins A and B separately. c) PGLa and different mutants of MAG2 [50]. See Table 2 for more details.

In another example, we consider AMPs in the Temporin family, namely Temporins A, B, and L [70]. Microbroth dilution assays to measure the inhibition of bacterial growth showed that against *E. coli* Temporin L was synergistic with Temporins A and B separately, while Temporins A and B were not synergistic. As shown in Fig. 7b, the Moran hydrophobicity autocorrelation is closer to each other for the Temporins A and B, and both of them exhibit relatively weak amphipathicity, as indicated by the slightly negative correlations at *d* = 9. However, there is a larger difference between the hydrophobicity autocorrelation parameters for Temporin L and the other Temporins A and B. The larger negative correlation reflects the greater amphipathicity for Temporin L compared to the other Temporins. These results suggest that the synergy between Temporin L and the other Temporins could be related to the differences in amphipathicity, while the similar levels of amphipathicity between Temporins A and B lead to the lack of synergy.

In the final example, we consider the peptide Magainin-2, originating from the Xenopus laevis frog, and several synthetic single-amino-acid-substituted analogs of Magainin-2 that are all synergistic with the peptide PGLA in killing *E*.*coli* bacteria [50]. Fig. 7c presents the Moran autocorrelation parameters for two distances *d* for all Magainin-2 species and for the PGLA peptide. One can see that autocorrelation parameters for PGLA are very different from all other Magainin-2 species. PGLA is non-amphipathic, while the negative correlations of different Magainin-2 peptides demonstrate strong amphipathicity. Conversely, it can be predicted that when the Magainin-2 peptides are combined in pairs, none of the pairs will be synergistic against *E. coli*. This is because the Magainin-2 peptides are all strongly amphipathic and thus too similar in this important physico-chemical characteristic. Therefore, PGLA and all considered Magainin-2 peptides are very different physicochemically, and this explains the observed antibacterial cooperativity in this system.

## Discussion

Recent experimental studies revealed that bacteria are more susceptible to combinations of some specific types of antimicrobial peptides. In this work, we present a theoretical investigation that allows us to identify synergistic pairs of AMPs based on their physicochemical properties. It is proposed that cooperating AMPs are those peptides that are the most different in their physical-chemical properties relevant to bacterial elimination. In other words, the more dissimilar are AMPs, the more cooperative they are in their anti-bacterial action. To test our hypothesis, we developed a computational framework that allowed us to quantify the physicochemical similarity and analyze its correlations with cooperativity in antibacterial activities. To illustrate our theoretical method, the synergy of AMP-AMP combinations acting against three different types of bacteria, namely *E. coli, M. luteus*, and *P. aeruginosa*, has been specifically considered.

A concept of physicochemical similarity between two peptides as inverse Euclidean distance in the space of properly normalized physicochemical features has been introduced and discussed. It has been found that there is a relatively small number of properties that are most relevant for supporting the synergy of AMP-AMP combinations. Theoretical analysis shows that measuring similarity using only the selected features inversely correlates with the antibacterial efficiency of AMP pairs, allowing us to separate synergistic and non-synergistic AMPs combinations. These observations clearly support our hypothesis that the most physicochemically dissimilar (in terms of the most relevant features) AMP pairs lead to cooperativity in the removal of bacterial infections, while similar AMP combinations do not exhibit cooperativity at all.

Generally, the selected physicochemical features for different bacteria do not coincide, although there are some common properties. It was found that several autocorrelation functions play an important role in supporting the synergistic action of AMPs combinations. Calculating these properties allowed us to explicitly illustrate the correlations between the physicochemical similarity and the antibacterial efficiency. In the considered several examples of AMP combinations acting against *E*.*coli* bacteria, cooperativity was observed only for peptides with very different autocorrelation parameters, while the peptides with similar autocorrelation parameters never produced synergistic combinations. This result again is fully consistent with our hypothesis about the relation between physicochemical similarity and antibacterial efficiency.

Although our theoretical approach connects the selected physicochemical properties of AMPs with their ability to cooperate, it does not provide a microscopic picture of underlying processes that lead to antibacterial synergy. At the same time, some of the obtained results allow us to present some speculations about possible molecular mechanisms of AMPs cooperativity. Our main idea is that this is a result of *complementarity* in antimicrobial properties of peptides. The possible microscopic picture is that one type of AMPs is affecting the bacterial membrane in a such way that it makes it easier for the second type of AMPs to disrupt the membrane. Since both activities are, in some sense, “orthogonal” to each other this lead to stronger effective cooperativity. Those AMPs that are similar in their physicochemical properties do not exhibit complementarity because in this case, peptides compete with each for the same regions of bacterial cellular membranes in order to disrupt them. One can also present these arguments as an effective free-energy landscape where similarity plays a role of the reaction coordinate as shown in Fig. 8. The pair of identical AMPs (A-A combination) or similar combinations (A-B Non-synergistic) do not have enough free energy to disrupt the bacterial cell, while dissimilar AMPs combinations (A-C synergistic) are able to affect bacteria because they have enough energy.

**Fig 8.**
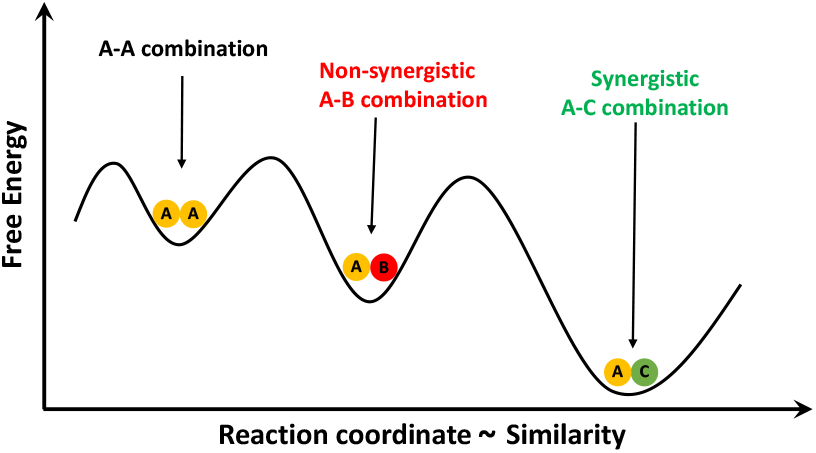
Free energy landscape description for AMP-AMP interaction.

Cooperativity is one of the main organizational principles in chemistry and biology required to support the functioning of living systems [71, 72]. Examples include multiple phenomena ranging from ligand binding to cellular receptors and enzyme activities to molecular machines made of complex protein complexes. In many cases, the molecular mechanisms of the processes that lead to cooperativity are still not fully understood. Although our theoretical approach provides a possible mechanistic explanation synergistic action of AMPs, it seems reasonable to suggest that similar ideas can also be extended to other biological processes. We propose that in some systems cooperating biological molecules might be complementary in their activities to perform their biological functions.

While our theoretical approach is successful in predicting the cooperativity of AMPs acting against bacteria, it is important to discuss its limitations. First, the method will work if there is enough data on synergistic and non-synergistic combinations for the given bacteria to identify the most relevant features and evaluate the physicochemical similarity. Second, it does not clarify the microscopic origin of cooperativity since it only detects the correlations but not their sources. Despite these limitations, however, this theoretical approach provides a powerful method for designing more efficient AMP drugs and it also gives the starting point for uncovering what molecular forces are responsible for the synergetic effects of peptides. In addition, this method can be easy to extend to other biological systems, e.g., to investigate cooperativity in protein-protein systems.

## Materials and methods

### Physicochemical similarity of AMP pairs

There are 1547 descriptors in total that are broadly divided into the following groups: autocorrelations (Moreau-Broto, Moran, and Geary coefficients for hydrophobicity, polarizability, free energy, and other features); amino acid compositions (single such as. for example, the percentage of valine in the peptide, or dipeptide, such as, for example, the percentage of valine adjacent to lysine); physicochemical compositions (composition and transition values for polarizability, charge, van der Waals forces, and other features), pseudo-amino acid compositions, and quasi-sequence order. Importantly, we view each peptide as a point in *d*-dimensional space of all physicochemical properties where each dimension represents a specific descriptor.

### Data normalization

Since the scales of physicochemical properties for each peptide are different, it is important as a first step to normalize them to have their values to be between 0 and 1. To do so, we subtract each descriptor value (*x*) from the corresponding mean (*μ*) and divide it by the standard deviation *σ*. The outcome is *z* = (*x* − *μ*)*/σ*. To normalize this quantity in the range 0 and 1, we use 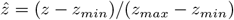.

### Euclidean distance as the measure of similarity between two peptides

There are different methods for defining the similarity between any two molecules [61, 62]. We chose the Euclidean distance in the space of all physicochemical descriptors so that the properties with greater distance would have greater weight in the calculation of overall distance. For each AMP-AMP combination, the Euclidean distance is calculated in terms of *N* different sets of descriptors,

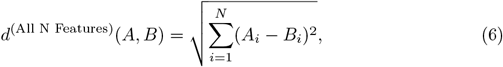

and for *M* selected features

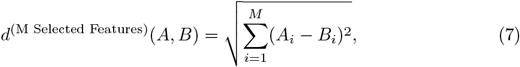

while, for individual descriptors we have

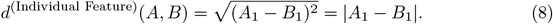

The Euclidean distance has been already successfully utilized in several bioinformatics investigations, including quantifying the relations between genes and proteins [73]. We postulate that the physicochemical similarity of two AMPs is inversely proportional to the Euclidean distance between them in the space of physicochemical properties,

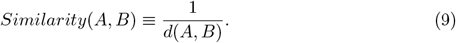

### Spearman’s rank correlation coefficient for calculating correlations between FIC values and Euclidean distance

The Spearman’s rank correlation coefficient is defined as the Pearson’s correlation coefficient between the ranks of distance, *R*(*d*), and ranks of FIC values, *R*(*FIC*),

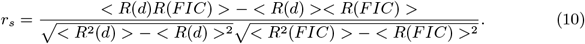

The Spearman’s rank correlation coefficient estimates how well the relationship between two variables can be described by a monotonic function, and, thus, it is a convenient measure to evaluate the correlations between two quantities. For computing Spearman’s correlation, one has to sort the values from least to greatest, and rank is the position of each sorted value in the list.

### Principal Component Analysis

PCA is a dimensionality reduction method that aims to preserve the local and global structure of the original data by applying a linear transformation [74]. The first principal component of a set of features *x*_*i*,1_, *x*_*i*,2_, …, *x*_*i,M*_ is the normalized linear combination of the features:

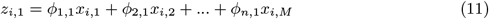

that has the largest variance subject to constraint that 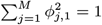. The corresponding Lagranian reads:

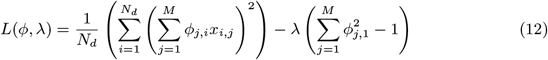

Where *N*_*d*_ is the number of data points. Solving this Lagrangian yields an eigenvalue equation:

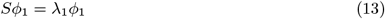

where *S* is the covariance matrix and *ϕ*_1_ = (*ϕ*_1,2_, *ϕ*_2,2_, …, *ϕ*_*M*,2_). Similarity we can find the second principal component *z*_*i*,2_ = *ϕ*_1,2_*x*_*i*,2_ + *ϕ*_2,2_*x*_*i*,2_ + … + *ϕ*_*M*,2_*x*_*i,M*_ by solving the corresponding eigenvalue equation *Sϕ*_2_ = *λ*_1_*ϕ*_2_.

Along the diagonal of the covariance matrix, large values point to patterns in the underlying structure of the data, and on either side of the diagonal, large values mean there are high correlations in the data and redundancies. Eigenvectors are averaged vectors

The variance represents the signal and the covariance represents redundancy. PCA aims to identify the direction in which the variance of the data is largest by searching orthogonally; these directions are principal components PCA assumes that the variance can be represented through linear transformations, that large variances represent the underlying structure of the data rather than noise, and that the principal components are orthogonal; the third assumption allows for a mathematically-convenient matrix composition so that the original matrix X can be represented as PX where P is the principal component.

The results of principal component analysis for all analyzed species are shown in Fig. 5, reflecting that the separation is strongest along the x-axis. The first component of PCA, along the x-axis, clearly separates synergistic and non-synergistic pairs, while the second component does not. It is possible that the pairs with strongest synergy show greater separation along the x and y dimensions, but future studies must test multiple AMP-AMP combinations on the same bacteria so that a greater range of FIC values is present. Lack of data and experimental error in the existing data could contribute to the lack of separation in the y-axis of the correlation and in the second component of PCA.

## Author Contributions

H.T. and A.M. designed the research. H.T. and A.M. performed the research. A.M. and H.T. and A.B.K. wrote the article.

## Data availability statement

The source code and data used to produce the results and analyses presented in this manuscript are available from Figshare repository: https://figshare.com/s/fa1408174d816c222b01.

## Acknowledgments

The work was supported by the Welch Foundation (C-1559), by the NSF (CHE-1953453 and CHE-2246878), and by the Center for Theoretical Biological Physics sponsored by the NSF (PHY-2019745).

